# The B.1.427/1.429 (epsilon) SARS-CoV-2 variants are more virulent than ancestral B.1 (614G) in Syrian hamsters

**DOI:** 10.1101/2021.08.25.457626

**Authors:** Timothy Carroll, Douglas Fox, Neeltje van Doremalen, Erin Ball, Mary Kate Morris, Alicia Sotomayor-Gonzalez, Venice Servellita, Arjun Rustagi, Claude Kwe Yinda, Linda Fritts, Julia Rebecca Port, Zhong-Min Ma, Myndi Holbrook, Jonathan Schulz, Catherine A. Blish, Carl Hanson, Charles Y. Chiu, Vincent Munster, Sarah Stanley, Christopher J. Miller

**Affiliations:** California National Primate Research Center, University of California Davis, Davis CA, USA; Center for Immunology and infectious Diseases, University of California Davis, Davis CA, USA; University of California, Berkeley, Department of Molecular and Cell Biology, Division of Immunology and Pathogenesis; Laboratory of Virology, Division of Intramural Research, National Institute of Allergy and Infectious Diseases, National Institutes of Health, Hamilton, MT, USA; Department of Pathology, Microbiology and Immunology, School of Veterinary Medicine, University of California Davis, Davis CA, USA; Division of Viral and Rickettsial Diseases, California Department of Public Health, Richmond, CA, USA; Department of Laboratory Medicine, University of California, San Francisco, San Francisco, CA, USA; Division of Infectious Diseases and Geographic Medicine, Department of Medicine, Stanford University School of Medicine; Division of Infectious Diseases, Department of Internal Medicine, School of Medicine, University of California Davis, Davis CA, USA

## Abstract

As novel SARS-CoV-2 variants continue to emerge, it is critical that their potential to cause severe disease and evade vaccine-induced immunity is rapidly assessed in humans and studied in animal models. In early January 2021, a novel variant of concern (VOC) designated B.1.429 comprising 2 lineages, B.1.427 and B.1.429, was originally detected in California (CA) and shown to enhance infectivity in vitro and decrease antibody neutralization by plasma from convalescent patients and vaccine recipients. Here we examine the virulence, transmissibility, and susceptibility to pre-existing immunity for B 1.427 and B 1.429 in the Syrian hamster model. We find that both strains exhibit enhanced virulence as measured by increased body weight loss compared to hamsters infected with ancestral B.1 (614G), with B.1.429 causing the most body weight loss among all 3 lineages. Faster dissemination from airways to parenchyma and more severe lung pathology at both early and late stages were also observed with B.1.429 infections relative to B.1. (614G) and B.1.427 infections. In addition, subgenomic viral RNA (sgRNA) levels were highest in oral swabs of hamsters infected with B.1.429, however sgRNA levels in lungs were similar in all three strains. This demonstrates that B.1.429 replicates to higher levels than ancestral B.1 (614G) or B.1.427 in the upper respiratory tract (URT) but not in the lungs. In multi-virus in-vivo competition experiments, we found that epsilon (B.1.427/B.1.429) and gamma (P.1) dramatically outcompete alpha (B.1.1.7), beta (B.1.351) and zeta (P.2) in the lungs. In the URT gamma, and epsilon dominate, but the highly infectious alpha variant also maintains a moderate size niche. We did not observe significant differences in airborne transmission efficiency among the B.1.427, B.1.429 and ancestral B.1 (614G) variants in hamsters. These results demonstrate enhanced virulence and high relative fitness of the epsilon (B.1.427/B.1.429) variant in Syrian hamsters compared to an ancestral B.1 (614G) strain.

**Author Summary:** In the last 12 months new variants of SARS-CoV-2 have arisen in the UK, South Africa, Brazil, India, and California. New SARS-CoV-2 variants will continue to emerge for the foreseeable future in the human population and the potential for these new variants to produce severe disease and evade vaccines needs to be understood. In this study, we used the hamster model to determine the epsilon (B.1.427/429) SARS-CoV-2 strains that emerged in California in late 2020 cause more severe disease and infected hamsters have higher viral loads in the upper respiratory tract compared to the prior B.1 (614G) strain. These findings are consistent with human clinical data and help explain the emergence and rapid spread of this strain in early 2021.

## Introduction

With the identification in humans of at least five circulating variants of concern (VOC) in the last 12 months from the UK, South Africa, Brazil, India, and California [1–3], it is apparent that SARS-CoV-2 VOCs will continue to emerge for the foreseeable future in the human population. These new variants are replacing formerly dominant strains and sparking new COVID-19 outbreaks. To avoid another uncontrolled SARS-CoV-2 pandemic, an ongoing effort is needed to monitor, collect and analyze data on new SARS-CoV-2 variants, identify VOCs and determine their impact on the performance of COVID-19 diagnostics and the efficacy of available treatments and vaccines.

In early January 2021, a novel VOC with 2 lineages, designated B.1.427 and B.1.429 (epsilon variant), was detected in California (CA) [4]. Epidemiologic studies suggest that the variants originated in CA and were responsible for an increasing proportion of cases beginning in November until more than 50% of the cases in in the state were due to the variant in February 2021 [4]. This suggested that epsilon (B.1.427/B.1.429) VOC had a moderately increased transmission efficiency in the population relative to the previously dominant B.1 (614G). In-vitro studies demonstrated that the L452R spike mutation found in B.1.427/B.1.429 increases infectivity and decreases susceptibility to antibody neutralization, rendering these CA VOCs completely resistant to one therapeutic monoclonal antibody, bamlanivimab [4]. B.1.427/429 has also been shown to partially escape vaccine elicited polyclonal and monoclonal neutralizing antibodies, using a novel mechanism of immune evasion [5,6]. A 2-3-fold reduction in the plasma neutralizing antibody titers against B.1.427/B.1.429 compared to ancestral B.1 (614G) variants in recipients of the Moderna mRNA vaccine was observed [5]. This finding was confirmed by a recent report showing that neutralizing titers in plasma from Wu-1 based mRNA vaccinees or from recovered patients were 2- to 3.5-fold lower against the B.1.427/B.1.429 VOC relative to B.1 (614G) pseudoviruses [6]. Further, the L452R mutation reduced neutralizing activity in a subset of receptor binding domain-specific monoclonal (m)Abs tested, and the S13I and W152C mutations abrogated neutralization by all N-terminal domain-specific mAbs tested [6]. Taken together these data provide evidence that the epsilon (B.1.427/B.1.429) VOC partially evades the human immune response [5,6].

Syrian hamsters are a widely used small animal model to study the infectivity and virulence of clinical SARS-CoV-2 isolates [7–11]. Following intranasal inoculation, hamsters are infected with SARS-CoV-2 and develop moderate to severe lung pathology. SARS-CoV-2−infected hamsters mount neutralizing antibody responses and are protected against homologous and heterologous re-challenge with SARS-CoV-2 [10,12,13]. In this study, we used the hamster model to determine the relative fitness and transmissibility of the ancestral SARS-CoV-2 and the B.1.427/429 strains in hamsters, and whether hamsters previously infected with the ancestral B.1 (614G) are susceptible to acute reinfection with B.1.427 or B.1.429.

We find that the B.1.427/429 strains are more virulent than the ancestral B.1 (614G) strain, as measured by weight loss of infected animals, viral titers in the upper respiratory tract and histopathology of the lungs. These findings are consistent with human clinical data and help explain the emergence and rapid spread of this strain in early 2021.

## Results

### Body weight loss in hamsters inoculated with SARS-CoV-2 epsilon (B.1.429/427) is more severe and sustained than in hamsters infected with ancestral B.1 (614G) SARS-CoV-2

To assess the virulence of SARS-CoV-2 strains, we infected hamsters intranasally with approximately 5000 PFU of B.1 (614G, B.1.427), or B.1.429 (Table 1). Body weights and oral swabs were obtained daily, and lungs collected at necropsy on 2, 4, 6 and 10 days post-inoculation were examined for pathologic changes and to determine the extent and level of virus replication (Table 1). Hamsters inoculated with SARS-CoV-2 began losing weight at day 2-3 PI with a nadir lasting from 4-6 days PI in B.1 (614G) animals, from 4-7 days PI in B.1.427 animals and from 4-8 days PI in B.1.429 animals. The difference in weight loss between the B.1 (614G) animals and epsilon (427/429) animals was statistically significant (Figure 1A).

**Figure 1.**
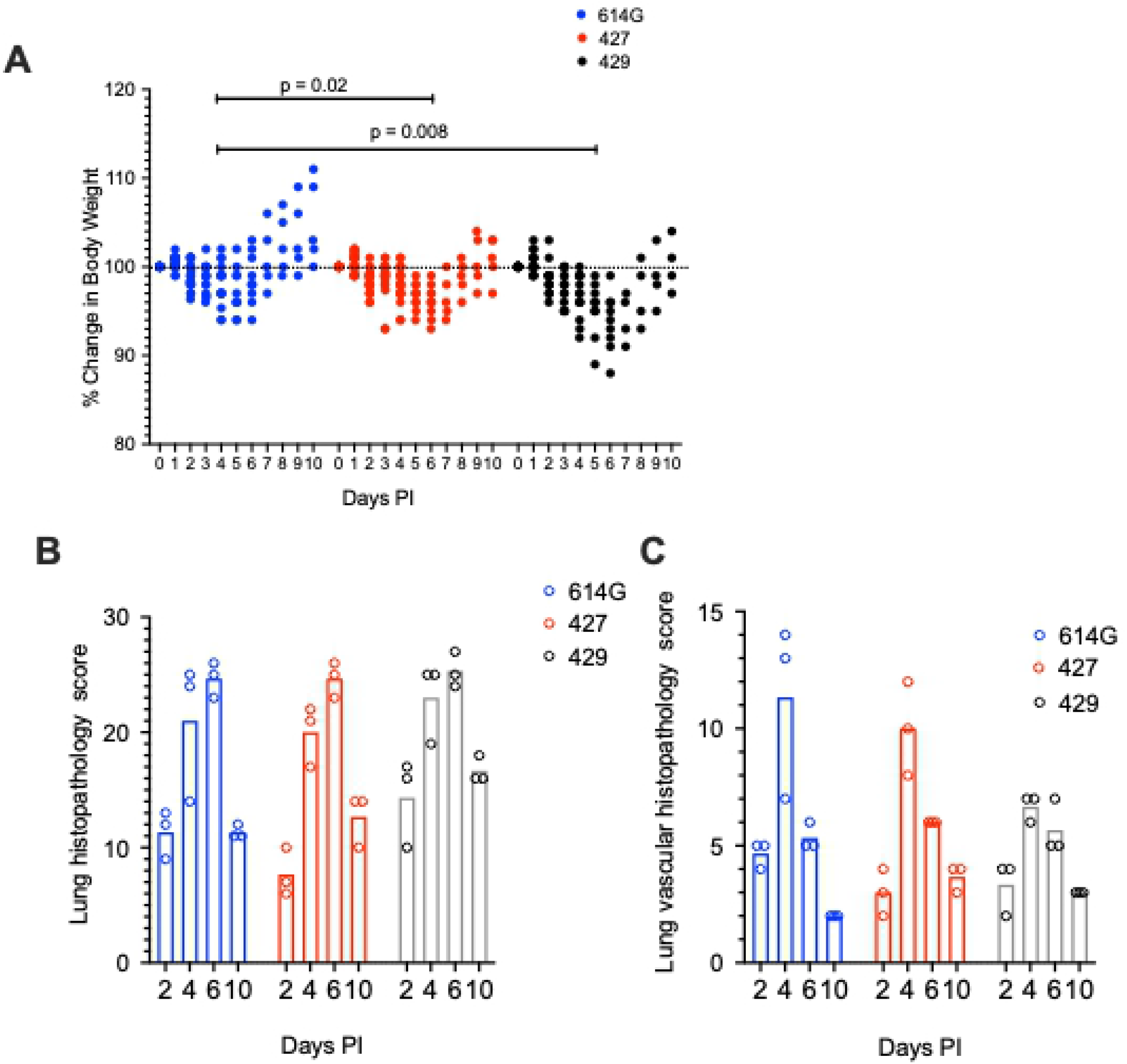
Change in body weight and lung histopathology scores in hamsters after intranasal inoculation with B.1 (614G), B.1.427 and B.1.429. **A)** Change in body weight relative to the day of inoculation. A mixed effects model was used to compare the groups. **B)** Total lung histopathology score (see methods for explanation). **C)** Lung vascular histopathology score (see methods for explanation). In A, mean values from each group were compared by fitting a mixed model, and a post-hoc multiple comparison test was used to compare the mean values of the 427 or 429 groups individually to the 614G group. Top of bars indicate mean values in B and C. .

**Table 1:** Summary of the experiments.

| Experiment | virus dose (PFU) | Virus | Oral swabs (Day PI) | Necropsy time points and animal numbers (Day PI/hamster number) | Figures |
| --- | --- | --- | --- | --- | --- |
| Pathogenesis/virulence | 5000 | B.1(614G) | d1, d2, d3, d4 | d2/6, d4/6, d6/6, d10/5 | 1, 2, 3, 4, 5 |
| Pathogenesis/virulence | 5000 | B.1.427 | d1, d2, d3, d4 | d2/6, d4/9, d6/6, d10/6 | 1, 2, 3, 4, 5 |
| Pathogenesis/virulence | 5000 | B.1.429 | d1, d2, d3, d4 | d2/4, d4/5, d6/5, d10/4 | 1, 2, 3, 4, 5 |
| Fitness/competition | 5000 | B.1.(614G) /B.1.427 - 1:1 | d1, d2, d3, d4 | d2/3, d4/3 | 6 |
| Fitness/competition | 5000 | B.1.(614G) /B.1.427 - 1:9 | d1, d2, d3, d4 | d2/3, d4/3 | 6 |
| Fitness/competition | 5000 | B.1.(614G)/B.1.427/ B.1.429/P.1/P.2/B.1. 1.1.7/ B.1.351 | d1, d2, d3, d4 | d2/5, d4/5 | 6, 7 |
| Prior infection/cross protection | 5000 | B.1(614G) then at 21 days PI B.1.(614G) | - | d2/5, d21/5, d23/5 | 8 |
| Prior infection/cross protection | 5000 | B.1.427 t then at 21 days PI B.1.427 | - | d2/5, d21/5, d23/5 | 8 |
| Prior infection/cross protection | 5000 | B.1(614G) then at 21 days PI B.1.427 | - | d23/5 | 8 |
| Prior infection/cross protection | 5000 | B.1(614G) then at 21 days PI B.1.429 | - | d23/5 | 8 |
| Airborne transmission | 10<sup>4</sup> | WA-1 | donors: d1<br>sentinels: DPE 0.5 -14 | donors: d 7 or 8/8<br>sentinels: up to DPE 14/8 | 9 |
| Airborne transmission | 10<sup>4</sup> | B.1.(614G) | donors: d1<br>sentinels: DPE 0.5 -14 | donors: d 7 or 8/8<br>sentinels: up to DPE 14/8 | 9 |
| Airborne transmission | 10<sup>4</sup> | B.1.427 | donors: d1<br>sentinels: DPE 0.5-14 | donors: d 7 or 8/8<br>sentinels: up to DPE 14/8 | 9 |
| Airborne transmission | 10<sup>4</sup> | B.1.429 | donors: d1<br>sentinels: DPE 0.5 -14 | donors: d 7 or 8/8<br>sentinels: up to DPE 14/8 | 9 |

### Intranasal inoculation of hamsters with SARS-CoV-2 B.1.429 results in more severe pulmonary pathology compared to B.1.427 or ancestral B.1 (614G) SARS-CoV-2

All hamsters inoculated with the SARS-CoV-2 variants developed moderate to severe broncho-interstitial pneumonia. To quantify the extent and severity of lung pathology, 2 scoring systems were used: the first evaluated all relevant changes in the lungs of the infected animals (Figure 1B) and the second evaluated only the pathology associated with the pulmonary vasculature (Figure 1C). In all animals, the extent of lung pathology increased from days 2-6 and decreased dramatically by 10 days post-infection (Figure 1B and C). However, the lungs from hamsters infected with B.1.429 had a trend toward higher overall pathology scores compared to animals infected with B.1 (614G) or B.1.427 (Figure 1B). B.1.429 inoculated animals exhibited more widespread and severe lesions at 2 days PI than B.1 (614G) and B.1.427 inoculated animals. These early severe histopathological changes also persisted for a longer time in B.1.429 infections as seen by comparison of the scores at days 2 and 10 (Figure 1B). Although not clinically significant, hamsters infected with B.1.429 tended to have less vascular pathology than animals infected with B.1 (614G) or B.1.427 (Figure 1C). B1.429 infection induced moderate to severe lung pathology more quickly and for a longer duration than B 1. (614G) or B1.427.

Although the extent, severity and timing of the lesions differed, the overall nature of the broncho-interstitial pneumonia was similar in all animals. At day 2 PI, lesions were centered on large airways and ranged from mild bronchitis to patchy, moderate bronchiolitis, bronchiolar epithelial cell necrosis and rupture of bronchiolar wall with limited extension of a mixed inflammatory infiltrate (composed of neutrophils, macrophages, fewer lymphocytes and scattered multinucleated syncytial cells) into adjacent alveolar septa (Figures 2 and 3: B, G, and L). The affected pulmonary surface area in examined sections ranged from 2 to 20% (Figure 2 B, G, and L). Vasculitis characterized by perivascular cuffing, intramural inflammatory cells and endothelialitis (sub-and intra-endothelial inflammatory cell infiltration), was noted in association with all SARS-CoV-2 variants, although more prominently in animals inoculated with B.1 (614G) and B.1.427. As noted above, the B.1.429 inoculated animals exhibited much more widespread and severe lesions at day 2 PI than B.1 (614G) and B.1.427 inoculated animals.

**Figure 2.**
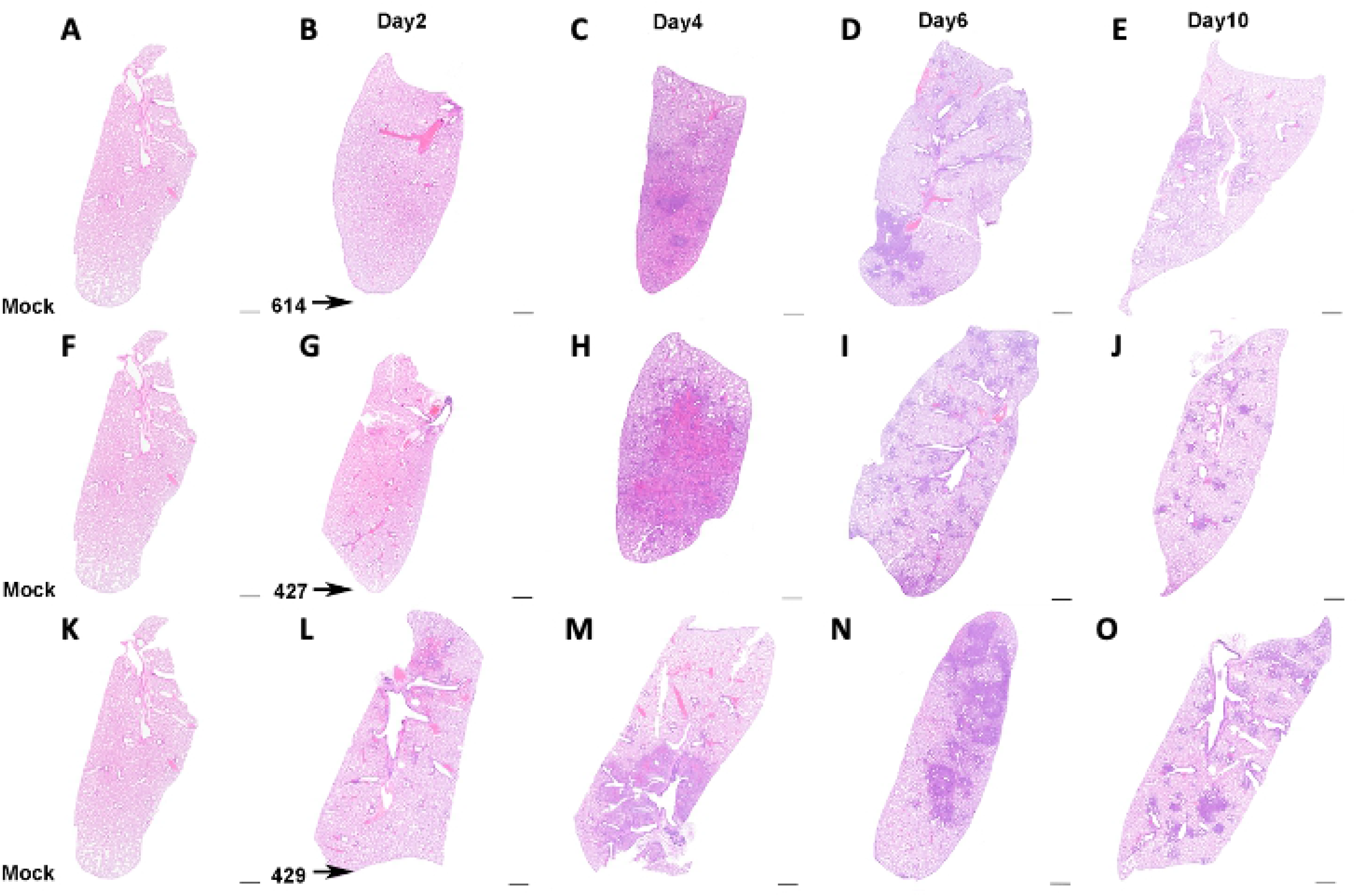
Extent of lung histopathology in hamsters after intranasal inoculation with B.1 (614G), B.1.427 and B.1.429. **A, F, K)** histology of normal lungs from uninfected hamsters. Top row: Histology of B.1 (614G) infection after **B)** 2 days PI, **C)** 4 days PI, **D)** 6 days PI, **E)** 10 days PI. Middle row: Histology of B.1.427 infection after **G)** 2 days PI, **H)** 4 days PI, **I)** 6 days PI, **J)** 10 days PI. Bottom row: Histology of B.1.429 infection after **L)** 2 days PI, **M)** 4 days PI, **N)** 6 days PI, **O)** 10 days PI. Hematoxylin and eosin staining. Sub-gross magnification. Scale bars equal 1 mm.

**Figure 3.**
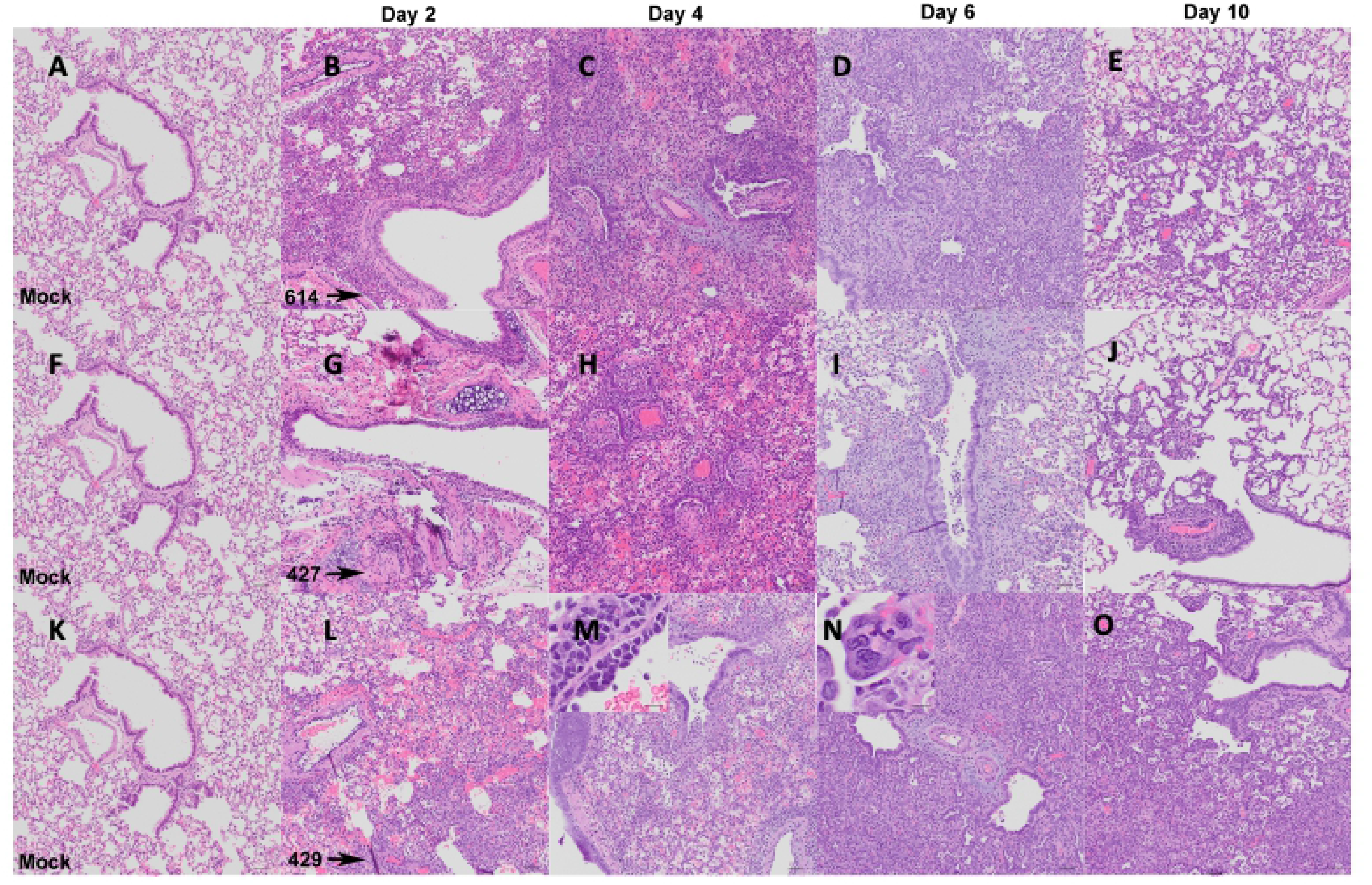
Nature of lung histopathology in hamsters after intranasal inoculation with B.1 (614G), B.1.427 and B.1.429. **A, F, K)** histology of normal lungs from uninfected hamsters. Top row: Histology of B.1 (614G) infection after **B)** 2 days PI, **C)** 4 days PI, **D)** 6 days PI, **E)** 10 days PI. Middle row: Histology of B.1.427 infection after **G)** 2 days PI, **H)** 4 days PI, **I)** 6 days PI, **J)** 10 days PI. Bottom row: Histology of B.1.429 infection after **L)** 2 days PI, **M)** 4 days PI, **N)** 6 days PI, **O)** 10 days PI. Hematoxylin and eosin staining. 100x magnification. Scale bars equal 50 um, inset scale bars equal 20 um.

By day 4 PI bronchiolar and alveolar lesions had progressed to necro-suppurative bronchiolitis with loss of normal alveolar septal architecture and replacement by hemorrhage, edema, fibrin, necrotic debris, mixed inflammation, and frequent multinucleated syncytial cells (Figure 2 and 3: C, H, and M), affecting up to 50% of the pulmonary surface area (Figure 2 C, H, and M). Perivascular cuffing and endothelialitis (vasculitis) remained prominent features, with mononuclear cells frequently extending from endothelium to adventitia in many small arteries (Figure 3 M, inset). Variable bronchiolar epithelial hyperplasia (characterized by epithelial cell piling up and increased mitotic figures) and scattered type II pneumocyte hyperplasia were also noted, particularly in those animals inoculated with B.1.429 (Figure 3 M).

Similar microscopic features, including necrotizing neutrophilic and histiocytic broncho-interstitial pneumonia with syncytial cells, perivascular cuffing, and endothelialitis, were observed at day 6 PI (Figures 2 and 3: D, I, and N), affecting 25-50% of the pulmonary surface area in examined sections (Figure 2 D, I, and N). Reparative changes, including bronchiolar epithelial hyperplasia and type II pneumocyte hyperplasia, were also prevalent at day 6 PI (Figures 2 and 3: D, I, and N).

By day 10 PI, the lung pathology in all animals had dramatically decreased. In animals inoculated with B.1 (614G) and B.1.427, necrosis, neutrophilic inflammation and vascular lesions appeared largely resolved, with replacement by patchy foci of mononuclear alveolar septal inflammation with bronchiolar epithelial and type II pneumocyte hyperplasia (Figures 2 and 3: E, J, O). The affected surface area in the lungs of these animals ranged from 2-10% (Figure 2 E, J, O). The animals inoculated with the B.1.429 variant exhibited more severe and widespread pulmonary lesions at day 10 PI (Figure 3 O), with persistence of neutrophilic to histiocytic alveolar septal inflammation with scattered syncytial cells and occasional endothelialitis as well as reparative changes. The affected surface area ranged from 15-20% in this group (Figure 2 O).

### Intranasal inoculation of hamsters with B.1.427 results in similar distribution of virus in lungs compared to ancestral B.1 (614G) SARS-CoV-2, while B.1.429 infects the lung parenchyma more rapidly

We used in-situ hybridization (ISH) to localize viral RNA (vRNA) to specific structures and cell types in the lung (Figure 4). The findings were similar in animals infected with either B.1 (614G) or B.1.427. At day 2 PI, bronchiolar epithelial cells were intensely labelled with infected cells extending along the entire length of main stem bronchi and smaller airways (Figure 4 A, E). In addition, rare focal areas of vRNA positive pneumocytes were found in the lung parenchyma. At day 4 PI in B.1 (614G) and B.1.427 animals, airway epithelial cells remained intensely labelled, with many of the infected cells detached from the basal lamina (Figure 4 B, F). However, most of the vRNA positive cells at day 4 were now found in the lung parenchyma, with type I and II pneumocytes and alveolar macrophages strongly positive (Figure 4B, F). In contrast, in the B.1.429 infected animals intense labeling of vRNA positive cells in the parenchyma and airways was already present at day 2 PI, and this persisted to day 4 PI (Figure 4 I, J). Of the 5-6 animals infected with each variant that were necropsied at day 6 PI, one animal from each group had a few vRNA positive cells in isolated foci in the lung parenchyma. The lungs from the remaining 4-5 animals in each group were negative (Figure 4 C, G, and K). At day 10 PI, vRNA+ cells were not found in the lungs of any of the animals (Figure 4 D, H, and L). Although B.1.429 disseminated to lung parenchyma more rapidly than B.1.427 or B.1 (614G), all 3 strains seemed to infect the same populations of cells in the lung: mainly airway epithelial cells and type I and type II pneumocytes (Figure 4). Alveolar macrophages were also labeled but this was likely to due to phagocytosis of infected cell debris rather than productive infection.

**Figure 4.**
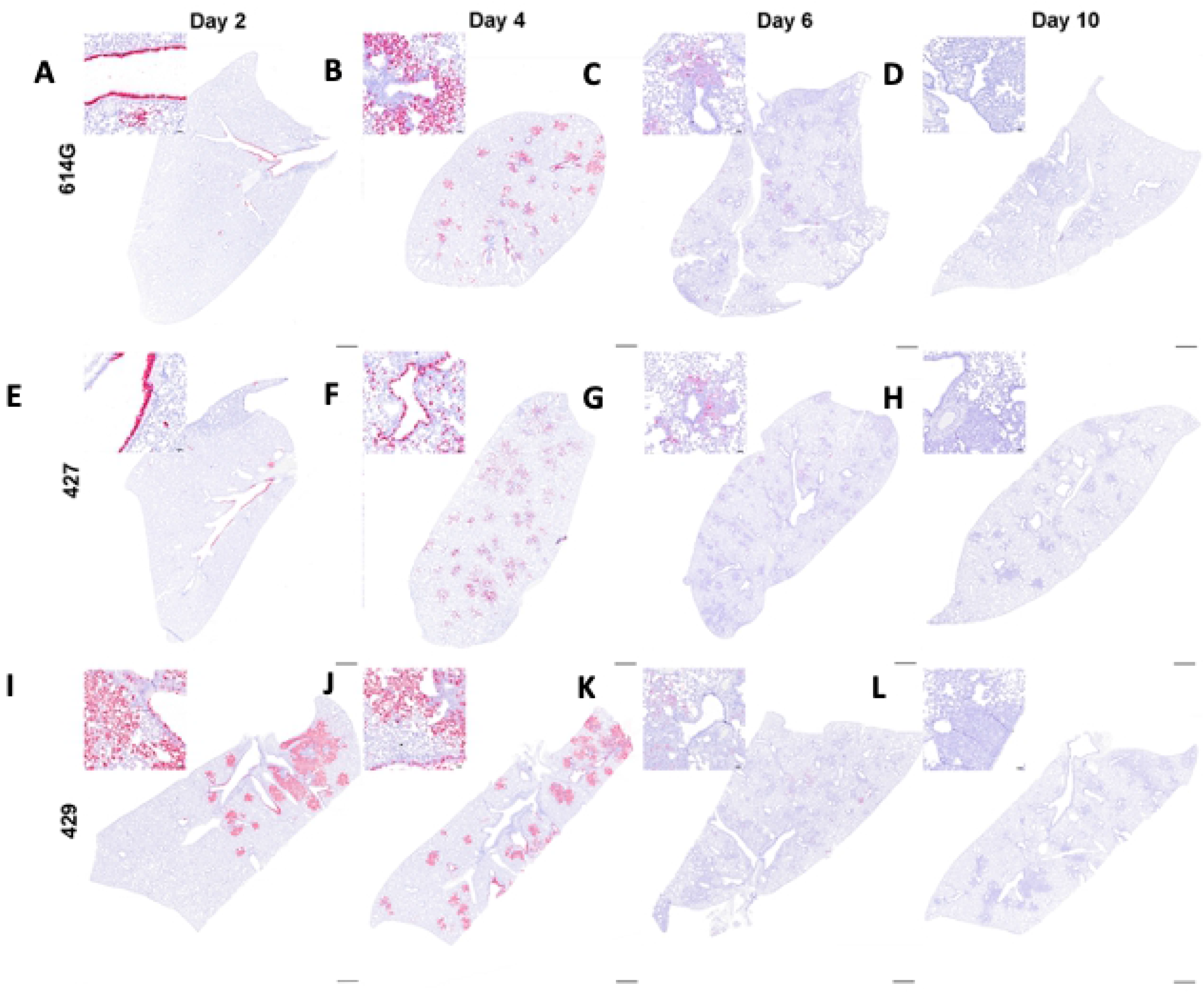
Distribution of SARS-CoV-2 RNA+ cells in lungs of hamsters after intranasal inoculation with B.1 (614G), B.1.427 and B.1.429 by in-situ hybridization (ISH). Sub-gross histology and magnified regions (inset) of ISH-labeled lung sections from hamsters infected with B.1 (614G) for **A)** 2 days PI, **B)** 4 days PI, **C)** 6 days PI and **D)** 10 days PI; or B.1.427 for **E)** 2 days PI, **F)** 4 days PI, **G)** 6 days PI and **H)** 10 days PI; or B.1.429 for **I)** 2 days PI, **J)** 4 days PI, **K)** 6 days PI and **L)** 10 days PI. Cells labeled by riboprobe in-situ hybridization stain red. Scale bars equal 1 mm.

### Intranasal inoculation of hamsters with B.1.429/427 results in similar virus kinetics and viral loads in lung and upper respiratory tract washes but sgRNA levels in oral swabs from B.1.429 infected animals were higher compared animals infected with to ancestral B.1 (614G) SARS-CoV-2

In all the SARS-CoV-2 infected hamsters, the levels of sgRNA in daily oral swabs were highest at 1 or 2 days PI and then declined steadily to day 4 PI. However, the sgRNA levels were significantly higher in the oral swabs of B.1.429 animals compared to the B.1.427 or B.1 (614G) animals (Figure 5A). The levels of sgRNA in URT washes collected at necropsy were highest at day 2 PI but had declined dramatically by day 10 PI (Figure 5B) in all animals and sgRNA levels were similar in all hamster groups (Figure 5B). The levels of infectious virus and sgRNA in the lungs of hamsters inoculated with all 3 viruses were very similar: high at days 2 and 4 PI but undetectable infectious virus and low sgDNA at days 6 and 10 PI (Figures 5C and D).

**Figure 5.**
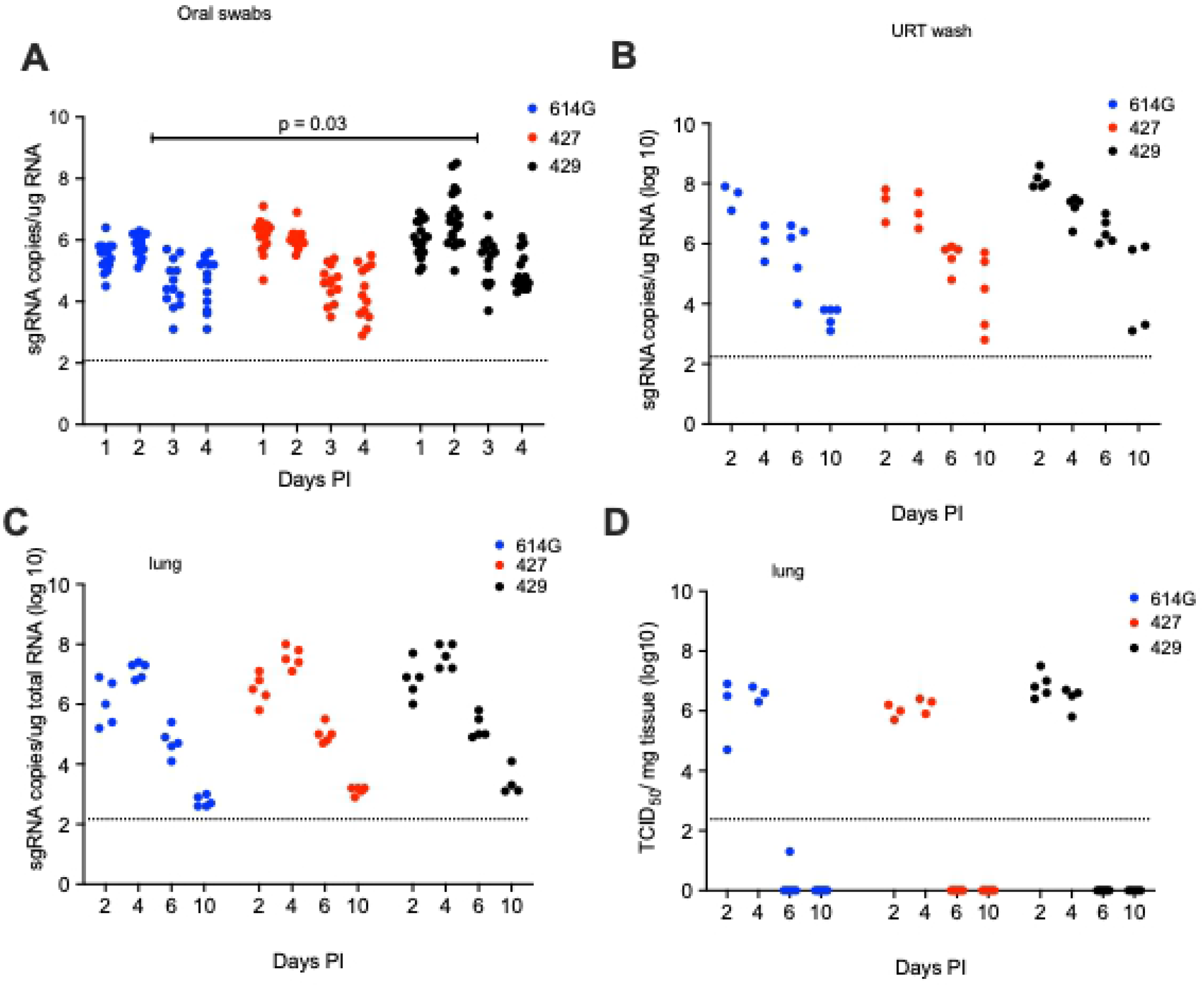
Viral loads in hamsters after intranasal inoculation with B.1 (614G), B.1.427 or B.1.429. **A)** sg RNA copies in oral swabs collected daily until day 4 or necropsy. **B)** sgRNA copies in upper respiratory tract (URT) washes collected at necropsy on 2, 4, 6 or days 10 PI. **D)** sgRNA copies in lungs collected at necropsy on 2, 4, 6 or days 10 PI. **D)** Infectious virus titers in lungs collected at necropsy on 2, 4, 6 or 10 days PI. In A, mean values from each group were compared by fitting a mixed model, and a post-hoc multiple comparison test was used to compare the mean values of the B.1.427 or B.1.429 groups individually to the 614G group.

### Specific variants predominate in lungs and nasal cavity of hamsters after intranasal inoculation with a mixture of SARS-CoV-2 variants

To determine if there is a relative fitness advantage among the circulating VOCs, hamsters were inoculated intranasally with a mixture of viruses and the proportion of each inoculated virus in the lungs and nasal cavity was determined. In the first experiment, animals (Table 1) were inoculated with 5000 PFU of SARS-CoV-2 that was either a 1:1 or 9: 1 mixture of B.1 (614G) and B.1.427 based on PFU. In hamsters inoculated with mixtures of 2 viruses, the level of sgRNA in the oral swabs were very highest at day 1 then declined until day 4 PI (Figure 6 A) while sgRNA levels in lungs and URT were higher at day 2 PI than day 4 PI (Figure 6 A, B and D). The sgRNA levels in the oral swabs, lungs and URT of hamsters infected with the 2 virus mixtures were very similar to animals inoculated with a single virus (Figure 6, Figure 5).

**Figure 6.**
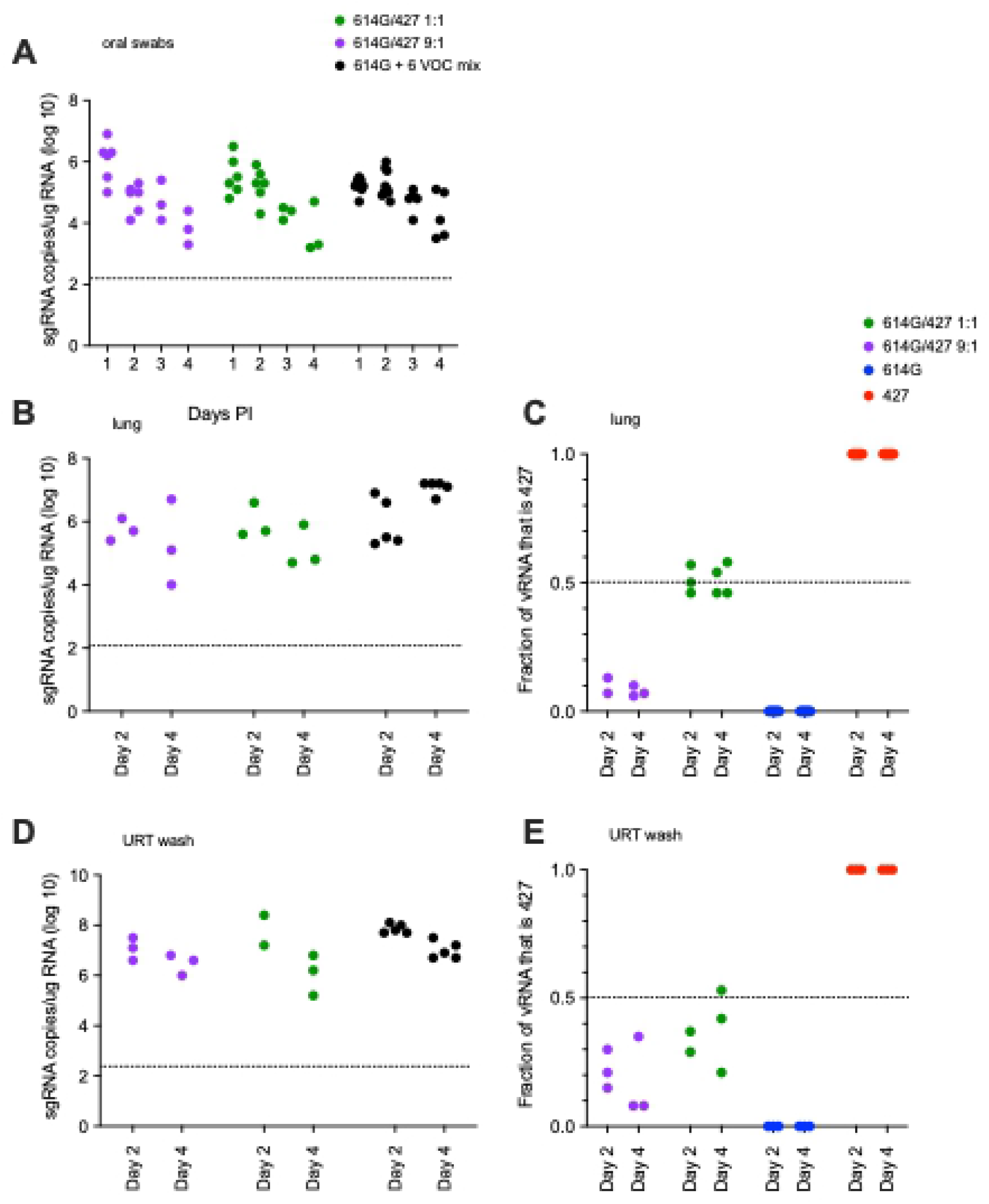
sgRNA levels and the proportion of each virus in hamsters after intranasal inoculation with a 1:1 or 9:1 mixed inoculum of B.1 (614G) and B.1.427; and sgRNA levels in hamsters inoculated with a mixed inoculum of 7 SARS-CoV-2 variants: B.1 (614G), B.1.427, B.1.427, P.1, P.2, B.1.1.7 and B.1.351. **A)** sgRNA copies in oral swabs collected daily for 4 days PI (copies/ug total RNA). **B)** sgRNA copies in lungs collected at necropsy on 2 and 4 days PI (copies/ug total RNA). **C)** proportion of the vRNA in lungs collected at necropsy on 2 and 4 days PI that was B.1.427. **D)** sgRNA copies in URT washes collected at necropsy on 2 and 4 days PI (copies/ug total RNA). **E)** proportion of the vRNA in URT washes collected at necropsy on 2, and 4 days PI that was B.1.427.

To determine if B.1 (614G) or B.1.427 had a competitive advantage over the other, RNA from the lungs and upper respiratory tract from the animals was sequenced to determine the proportion of each virus in the sample. In the URT and lung samples of the 1:1 inoculated animals, B.1.427 made up between 39-57% of the virus population at day 2 PI and 21-58% at day 4 PI (Figure 6C and E). In the lung samples of the 9:1 inoculated animals, B.1.427 made up about 10% of the virus population in the lungs at day 2 and 4 PI (Figure 6C), while in the URT, B.1.427 made up from 8% to 35% of the virus population at day 2 and 4 PI (Figure 6E). These results suggest that B.1.427 may have slight replicative advantage in the URT compared to B.1 (614G). However, there was no indication that B.1.427 had an enhanced ability to replicate in lungs compared to B.1 (614G).

To simultaneously determine the relative fitness of a larger number of variants, in a third experiment, 10 animals (Table 1) were inoculated with 5000 PFU of SARS-CoV-2 that was composed of an equal mixture, based on PFU, of 7 SARS-CoV-2 variants: ancestral B.1 (614G) [18], B.1.427 [4], B.1.429[4], P.1 [19], P.2 [19], B.1.1.7 [20] and B.1.351 [21]. We developed a amplicon sequencing strategy named QUILLS (<u>QU</u>asispecies Identification of <u>L</u>ow-<u>F</u>requency <u>Li</u>neages of SARS-CoV-2) to identify relative frequencies of viral variants within a mixed population by sequencing of key single nucleotide mutations in the spike and orf1ab genes (see Methods). QUILLS analysis of the pure viral stocks used to generate the mixture revealed > 99.5% of the RNA sequences obtained from the stocks were identical to the published sequence of the respective VOC; analysis of mixture showed that each VOC comprised between 4% and 25% of the RNA in the mixed virus inoculum (Figure 7A), as expected based on PFU normalization. In the hamsters inoculated with this mixture of 7 viruses, the levels of sgRNA in the lungs and URT were very high at days 2 and 4 PI (Figure 6 B and D) and were similar to the levels found in animals inoculated with a single virus (Figure 5 C and D).

**Figure 7.**
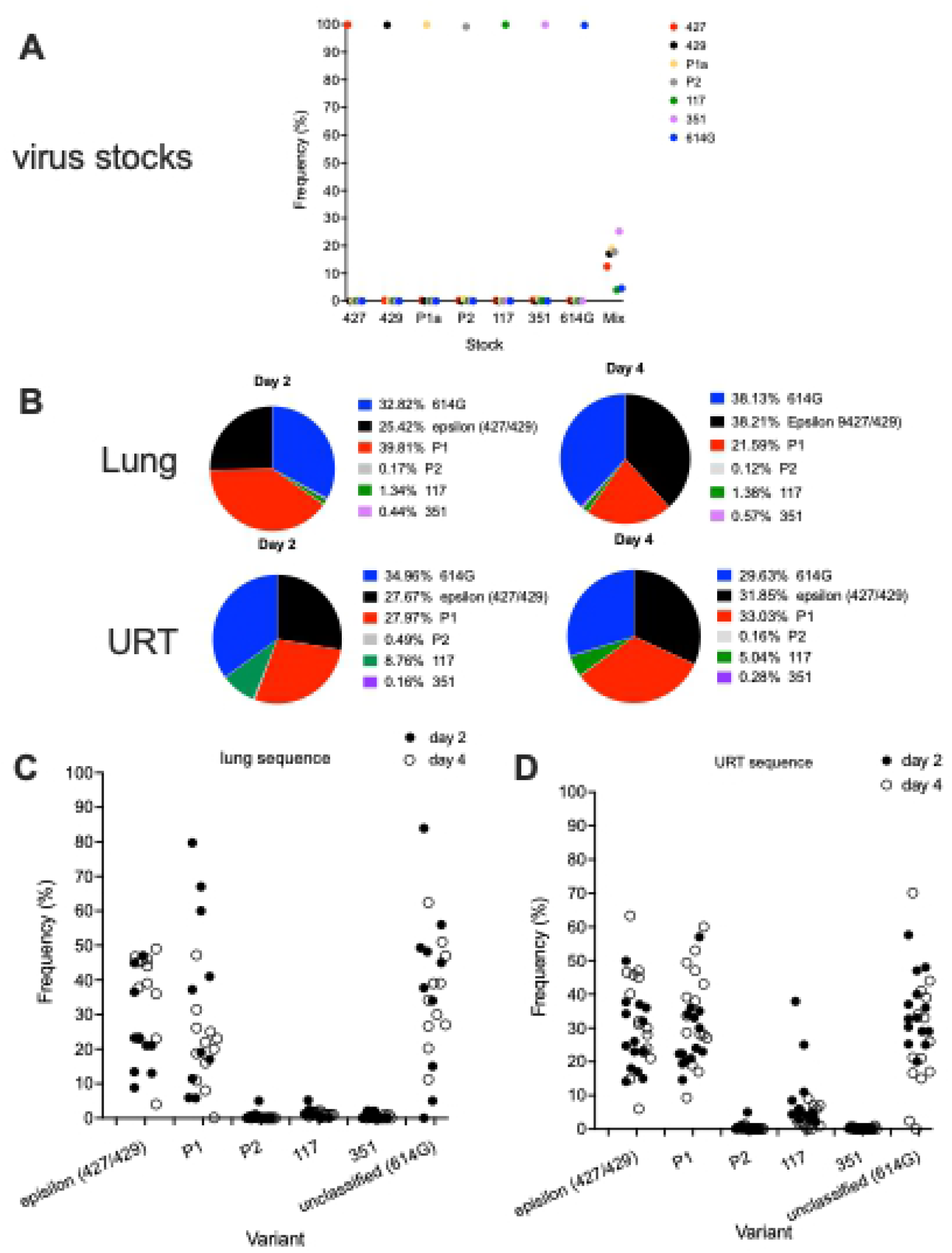
Relative levels of SARS-CoV-2 variants in hamsters after intranasal inoculation with a mixed inoculum of 7 SARS-CoV-2 variants: B.1 (614G), B.1.427, B.1.429, P.1, P.2, B.1.1.7 and B.1.351. **A)** Proportion of each variant RNA in the total vRNA of each virus stock and the mixed inoculum. **B)** Mean proportion of each variant RNA in the total vRNA at 2 and 4 days PI in the lungs (upper row) and URT (lower row) of hamsters infected with the mixed inoculum. **C)** Proportion of each variant RNA in the total vRNA at 2 and 4 days PI in the lungs of each hamster infected with the mixed inoculum. **D)** Proportion of each variant RNA in the total vRNA at 2 and 4 days PI in the URT washes of each hamster infected with the mixed inoculum.

To determine if one or more of the 7 variants in the inoculum had a competitive advantage over the others, RNA from the lungs and upper respiratory tract from animals necropsied at day 2 PI (n=5) and day 4 P (n=5) was sequenced and the proportion of each virus in the samples was determined. In the day 2 lung samples, B.1 (614G), P.1 and epsilon (B.1.427/B.1.429) predominated (mean: 32.8%/ range: 0-84%, 38.8%/0-98% and 25.4%/1-45% respectively) with P.2, and 351 making up less than 2% of the vRNA (Figure 7 B and C). On day 4, the lung vRNA was B.1(614G): 38.1%/11.2-51%, epsilon (B.1.427/ B.1.429) 38.2%/4-49%, P.1: 21.6%/0.1-47.2%, with B.1.1.7, P.2 and B.1.351 making up less than 2% of the vRNA (Figure 7 B and C). Thus, in the lung, B.1 (614G), P1 and epsilon (B.1.427/ B.1.429) were the most frequent variants while B.1.1.7, B.1.351 and P.2 were only found at low levels.

In the day 2 PI URT washes, the vRNA was B.1 (614G): 35%/20-48%, P1: 28%/19.4-57%, epsilon (B.1.427/ B.1.429): 27.7%/14.1-49.9%, B.1.1.7: 8.8%/2-37.9%, with P2 and 351 making up less than 1% of the vRNA (Figure 7 B and D). On day 4, the URT vRNA was B.1 (614G): 29.6%/0-70.2%, P.1: 33%/9.7-60%, epsilon (B.1.427/ B.1.429): 31.9%/14-63.3%, B.1.1.7: 5%/1-13%, with P.2 and B.1.351 making up less than 1% of the vRNA (Figure 7 B and D). Thus, in the URT, B.1 (614G), epsilon (B.1.427/ B.1.429) and P.1 were the most frequent variants, B.1.1.7 was intermediate and B.1.351 and P.2 were infrequent. These results demonstrate that B.1, P.1 and epsilon (B.1.427/ B.1.429) have a competitive advantage over the other variants in the lung but that B.1.1.7 can compete with P.1 and epsilon (B.1.427/ B.1.429) in the URT.

### Prior Infection with B.1 (614G) protects hamsters from subsequent challenge with B.1.427/429

To confirm protection from homologous challenge, hamsters (Table 1) were inoculated intranasally with 5000 PFU B.1 (614G) and then were rechallenged with 5000 PFU B.1 (614G) 21 days later (Figure 8A); another group of hamsters (Table 1) was infected with B.1.427 and rechallenged with B.1.427 21 days later (Figure 8B). To determine if B.1.427 and B1.429 are susceptible to the immune responses elicited by prior infection with B.1 (614G), hamsters (Table 1) were inoculated intranasally with 5000 PFU of B.1 (614G) and then were challenged 21 days later by intranasal inoculation of 5000 PFU of B1.427 or B1.429. To document infection and to determine the levels of vRNA at time of challenge, five animals in each group were necropsied at day 2 and day 21 respectively (Table 1). Rechallenged animals were necropsied on day 2 days after rechallenge (day 23) (Table 1). Viral titer and vRNA levels in lungs were determined for all groups and timepoints (Figure 8).

**Figure 8.**
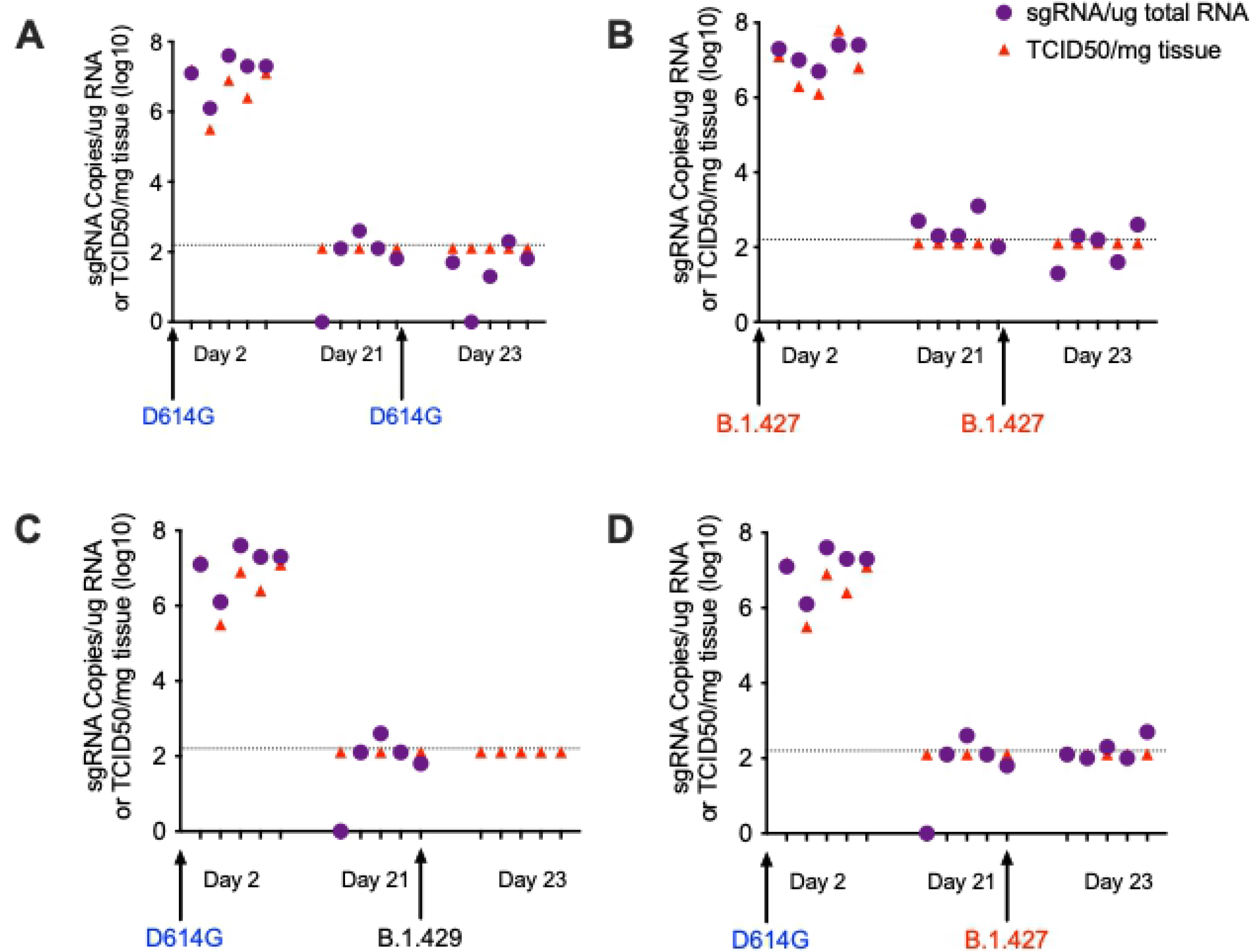
Prior infection with B.1 (614G) protects hamsters from subsequent challenge with B.1.427 or B.1.429. **A)** sgRNA levels and infectious virus titer in hamster infected with B.1 (614G) and then challenged 21 days later with homologous B.1 (614G). **B)** sgRNA levels and infectious virus titer in hamster infected with B.1.427 and then challenged 21 days later with homologous B.1.427. **C)** sgRNA levels and infectious virus titer in hamster infected with B.1 (614G) and then challenged 21 days later with heterologous B.1.429. **C)** sgRNA levels and infectious virus titer in hamster infected with B.1 (614G) and then challenged 21 days later with heterologous B.1.427.

All animals had high levels of sgRNA and infectious virus in lungs at day 2 PI, but no infectious virus and only very low levels of sgRNA were detected in animals at day 21 PI (Figure 8), confirming that the animals had been infected by the initial inoculum and then cleared the infection. Two days after re-challenge (day 23 PI), we could not isolate virus or detect sgRNA in the lungs of any of the animals (Figure 8). Thus, prior infection with ancestral B.1 (614G) protects hamsters from challenge 21 days later with either B.1.427 or B.1.429, with protection defined as no virus replication in lungs.

### Th efficiency of airborne transmission between hamsters infected with (B.1.427/429) and B.1 (614G) is similar

To determine if the airborne transmission of B.1.427 and B1.429 between hamsters is more efficient than B.1 (614G) airborne transmission, four groups of eight “donor” hamsters (Table 1) were inoculated intranasally with 10^4^ PFU of either B.1 (614G), A.1 (WA-1), B.1.427, B.1.429 (Figure 9). Oral swabs were collected at day 1 PI and naïve “sentinel” animals were added to one end of the cage separated from the donor animals by a barrier that prevents large particles but allows smaller particles to pass between the co-housed animals [22]. Donor animals were necropsied at Day 7 or 8 PI and “sentinel” animals were monitored for virus in oral swabs collected for up to 14 days after exposure. High levels of genomic and sgRNA were detected in the oral swabs of all the donor animals confirming that they were infected (Figure 9A). It is worth noting that the day 1 PI sgRNA levels in the swabs of B.1.429 infected animals were the highest and the difference compared to WA-1 infected animals was significant (Figure 9A). Of the eight sentinels exposed to the B.1.427 donors, all 8 were infected by 2 days post-exposure (PE). By comparison, 100% of B.1 (614G) sentinels were infected by 3 days PE and 100% of WA-1 sentinels were infected by 5 days PE (Figure 9B). Thus, B.1.427 was transmitted marginally more rapidly than WA-1 and B.1 (614G). In contrast among the B.1.429 sentinels, 7 out of 9 animals were infected by 3 days PE and 8 of 9 animals were infected by 5 days PE (Figure 9B). Thus, B.1.427 transmits between hamsters marginally more efficiently than B.1 (614G) and WA-1, while B.1.429 transmits marginally less efficiently than B.1 (614G) and WA-1. Taken together, these data suggest that airborne transmission of epsilon (B.1.427/429) and B.1 (614G) between hamsters is similar.

**Figure 9.**
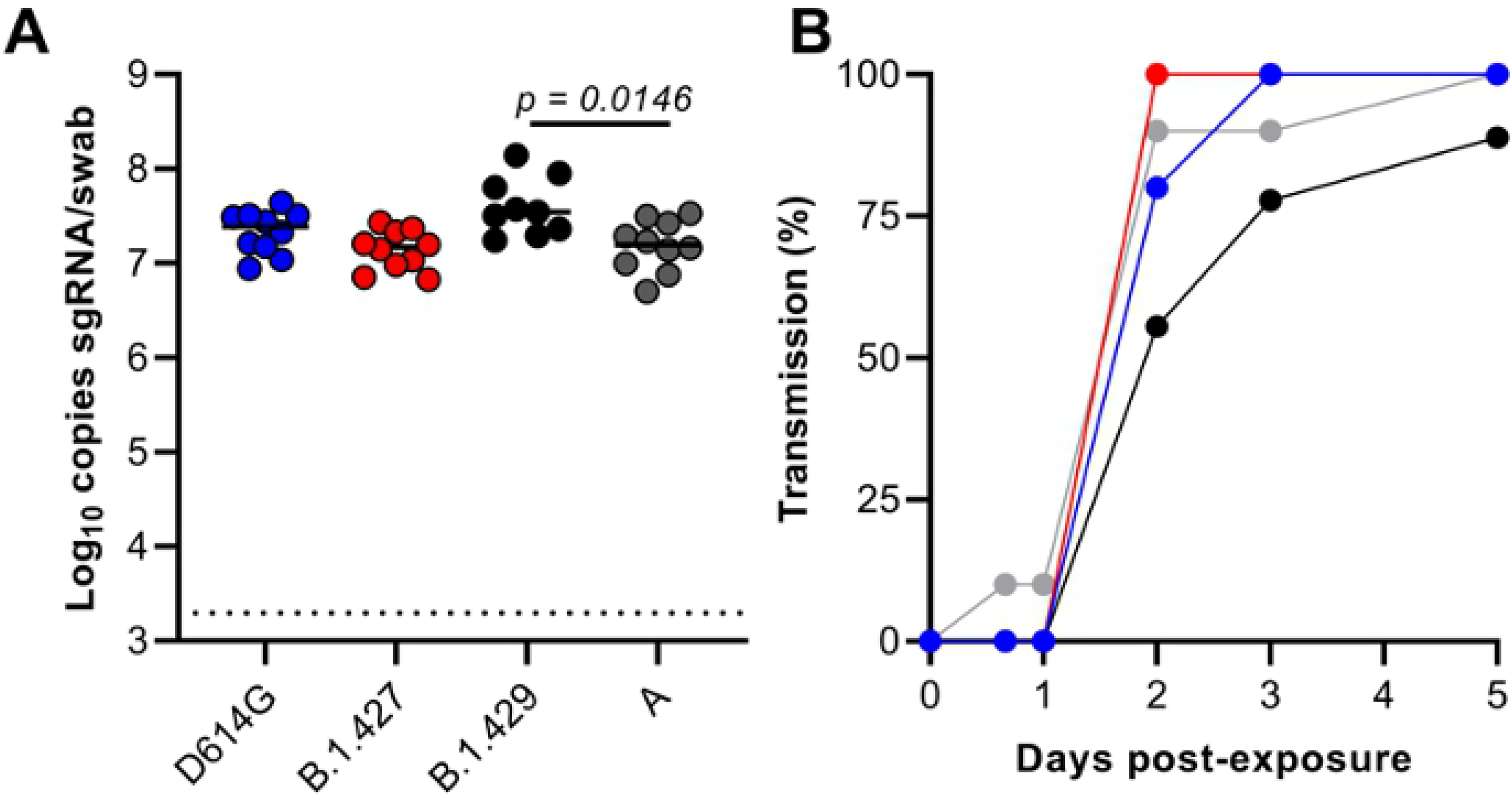
Relative efficiency of B.1 (614G), B.1.427 and B.1.429 airborne hamster to hamster transmission. **A)** sg RNA levels in oral swabs collected from donors 1 day after intranasal inoculation. **B)** percent of sentinel animals that are infected each day based on detection of sgRNA in oral swabs collected at least daily after co-housing.

## Discussion

The SARS-CoV-2 epsilon variant is comprised of 2 separate lineages, B.1.427 and B.1.429, with each lineage rising in parallel in California and other western states [4]. The variant is predicted to have emerged in California in May 2020 and increased in frequency from 0% to >50% from September 2020 to January 2021. The B.1.427/B.1.429 variant is no longer the predominant circulating strain in California, as it was replaced first by the B.1.1.7 (alpha) variant, which has since been replaced by the B.1.617.2 (delta) variant. The SARS-CoV-2 B.1.427/B.1.429 (epsilon) variant has a characteristic triad of spike protein mutations (S13I, W152C, and L452R) [4]. Epidemiologic and in-vitro studies found that the variant is approximately 20% more transmissible with 2-fold increased shedding in patients compared to ancestral B.1 (614G) [4] and that the spike L452R mutation conferring increased the infectivity of pseudoviruses in vitro [4]. It was not clear if the B.1.427/B.1.429 variants caused more severe disease; however, as the frequency of infection with the B.1.427/B.1.429 variants increased, the number of cases increased [4], followed by an increase in COVID-19 associated hospitalizations and death (https://covid19.ca.gov/state-dashboard/). In this study, we demonstrated that, based on differences in weight loss, SARS-CoV-2 epsilon (B.1.429 and B.1427) is more virulent than ancestral B.1 (614G) SARS-CoV-2 and B.1.429 is more virulent than B.1.427 in Syrian hamsters. The more rapid dissemination of virus from the airways to the alveoli in the lungs and higher levels of virus replication in the URT (daily oral swabs) of B.1.429 infected animals provide a biologically plausible explanation for the higher virulence of this L452R carrying strain.

The significantly higher sgRNA levels in the oral swabs collected daily over the first 4 days of B.1.429 infection suggests that this variant has enhanced fitness in the URT of hamsters compared to B.1 (614G) or B.1.427. This finding is consistent with the finding that there is a 2-fold increase in median viral loads in nasal swabs of patients infected with B.1.427/B.1.429 SARS-CoV-2 (epsilon) compared to patients infected with ancestral B.1 (614G) [4]. However, it is important to note that the overall levels of sgRNA in the lungs of animals infected with all three viruses were similar. Although not statistically significant, it should also be noted that the sgRNA levels in the B.1.429 infected animals were higher at all time points compared to levels in B.1 and B.1.427 infected animals (Figure 5B). The inability to detect a significant difference in sg RNA levels in URT washes collected at necropsy is likely due to the fact only 3-5 URT wash samples were collected from each group of hamsters on days 2 and 4 PI, while 16-19 oral swab samples were collected daily from each group of hamsters on days 1-4 PI. The URT-specific enhanced fitness of B.1.429 could be explained by a higher density of suspectable and permissive cells for B.1.429 in the URT compared to the lung or by the presence of mutations in B.1.429 that makes it more adapted for replication in the URT. Prior studies have suggested that the L452R mutation may increase infectivity because it stabilizes the interaction between the spike protein and its human ACE2 receptor [23] [24]. In-vitro studies found that the L452R mutation that defines the episilon VOC enhances pseudovirus infection of 293T cells and lung organoids [4]. Our finding in hamsters of higher replication levels of B.1.429 in the URT, but not the lung, compared to the ancestral B.1 (614G) variant extends these in-vitro findings to intact animals. Notably, our competition experiments demonstrated that the L452R containing B.1.427 and B.1.429 variants outcompeted many other variants that do not contain L452R (B.1.1.7, P.2, B.1.351) in the lungs and URT. The exceptions were B.1 (614G) and the P1 variant, which replicated to levels that were similar to (epsilon) B.1.427/B.1.429 in both lungs and URT; and the B.1.1.7 variant that was able to compete with B.1 (614G), B.1.427/429 and P.1 in the URT but not the lung. Our results are consistent with a previous report that the B.1.1.7 variant replicates to higher levels in the nose of Syrian hamsters than the ancestral B.1 variant [25]. Both the P.1 and B.1.1.7 variants carry the N501Y-mutation that also enhances infectivity in-vitro [24] and this may explain why they can successfully compete with the L452R variants in the nose of hamsters. It is also possible to conclude from these results that the relative fitness of different variants depends, at least to some extent, on the anatomic site of the infection.

The QUILLS strategy to identify relative frequencies of viral variants within a mixed population by sequencing of key single nucleotide mutations has limitations in that only variants with known mutations can be identified. All the variants used in the mixed inoculum experiments in this study, have lineage-defining mutations in spike and orf1ab on the backbone of ancestral B.1 (614G) and these 6 variants are unambiguously identified in the infected animals by QUILLS. However, the variant identified as B.1(614G) using QUILLS is an aggregate of all variants carrying 614G that cannot be otherwise be assigned to one of the other 6 variants in the inoculum. This would include novel variants arising from the inoculum B.1(614G) and revertants that may arise in each VOC quasispecies. Because of this, the relative frequency assigned to the ancestral B.1(614G) is less reliable than the frequencies of the other 6 variants that are based on positively identified sequences in the samples. Similarly, a single point mutation in orf1ab, that can arise or revert spontaneously, distinguishes B.1.427 and B.1.429. This makes it difficult to unambiguously identify B.1.429 and B.1.427 in mixtures and thus we reported the results as the frequency of the “epsilon”, the arrogate of the B.1.429 and B.1.427 frequencies.

As previously reported with ancestral B.1 and other VOC [11,25,26], in this study intranasal inoculation of hamsters with the SARS-CoV-2 B.1 (614G) or epsilon variants (B.1.247/B.1.249) induced progressive moderate to severe broncho-interstitial pneumonia with vasculitis, beginning as early as day 2 PI. By day 10 PI, lesions were largely resolved, with only residual mononuclear interstitial inflammation and reparative changes. While the pulmonary pathology in hamsters largely mirrored that of humans, unlike humans that die of COVID [14–17], we did not observe thrombosis, microangiopathy or necrotic vessel walls in any of the hamsters. However, all hamsters recover from the SARS-CoV-2 infection, and thus do not model fata COVID disease. This difference makes the comparison of lung pathology of hamsters and humans imperfect. While the nature and distribution of the lesions were similar with all strains, by analyzing histologic changes frequently (day 2, 4, 6 and 10 PI), we were able to detect differences in the timing of the onset and resolution of the lesions in animals infected with different strains. Although we did find that hamsters infected with B.1(614G) and B.1.427 tended to have more severe vascular changes than animals infected with B.1.429, this increased vascular disease was not associated with more weight loss or higher levels of viral replication and may be an incidental finding. We show here that studies employing sequential analysis of pathologic changes in larger numbers of inoculated hamsters can detect differences in the time course of disease between SARS-CoV strains. Studies that examine one or two timepoints PI are less likely to be informative. In fact, although a study comparing B.1.1.7 and B.1.351 infection in female Syrian hamsters found no major differences in lung histology, only one time point after infection, day 4 PI, was examined [26]. Even at this single time point however, compared to ancestral B.1 strains, the proinflammatory cytokine response was more intense in lungs of B.1.1.7 infected hamsters [26]. This discordance between the relative intensity of inflammation and the relative levels of inflammatory mediators reported demonstrates that sequential examination of both parameters is needed to determine how the lung inflammatory cell infiltrates drive the differential expression of inflammatory cytokines.

The ISH results demonstrated that B.1.427 and B.1 (614G) SARS-CoV-2 variants disseminate from the airways to the lung parenchyma at between days 2 and 4 PI, at least 24 hours after B.1.429 has begun replicating in alveoli. The faster observed dissemination of virus to alveoli in B.1.429 is consistent with the more rapid onset of histologic changes in the lung. The combination of ISH and histopathologic examination of lungs collected at frequent intervals post-inoculation seems to be a good strategy to detect differences in SARS-CoV-2 VOCs virulence.

An ongoing concern is the extent to which newly emerging variants can evade preexisting immunity that was generated from infections with ancestral SARS-CoV-2 strains. These concerns have been validated by the P.1 epidemics in Manaus, Brazil that occurred despite seropositivity rates of up to 76% prior to P.1 emergence [19]. The result of our serial infection experiments clearly demonstrate that immune responses present 21 days after primary infection by ancestral B.1 (614G) infection protect hamsters from challenge with both epsilon variants (B.1.427/B.1.429). In fact, we were unable to detect any virus replication, or infectious virus, in the lungs of any of the previously infected hamsters 2 days after the heterologous challenge. Thus, the immunity induced by the prior B.1 (614G) infection completely protected the lung from the epsilon infections and thus the animal from significant disease. Similarly, strong protection is found in hamsters infected with B.1 and rechallenged with B.1.1.7 [25]. We did not however assess the levels of virus replication in the URT after heterologous challenge, thus the degree to which prior infection affects replication in the URT, and therefore transmission between hamsters, is unknown. In addition, as the rechallenge occurred at day 21 PI, mature antiviral immune responses are expected to be present in blood and secretions [10,13], however, we did not make any attempt to identify the immune mechanism associated with the heterologous protection.

Epidemiologic studies led to the estimate that the epsilon (B.1.427/B.1.429) variant is 20% more transmissible than the ancestral B.1 (614G) and this was attributed to higher viral loads in the URT [4] . However, the hamster transmission studies reported here failed to show a consistent or large difference in transmission efficiency, between the epsilon (B.1.427/ B.1.429) variant and the ancestral B.1 (614G) strain, despite the higher viral load in the URT of B.1.429 infected hamsters. The inability to model increased airborne transmission of the epsilon (B.1.427/ B.1.429) variant was likely due to a combination of the experimental design and the relatively small differences in human transmission efficiency between B.1 (614G) and epsilon (B.1.427/ B.1.429). Differences in hamster transmission would likely be apparent with a variant that was at least 50% more transmissible than the ancestral strain and/or if the sentinel animals were exposed to donor animals when virus shedding is no longer at its peak, 2-3 days after they were inoculated. This may detect differences in transmission based on duration of URT shedding, while the 24 hour PI donor exposure approach attempts to detect differences due to the levels of URT shedding at the peak of virus replication.

As novel SARS-CoV-2 variants continue to emerge, it is critical that their pathogenic potential be rapidly assessed in animal models. We found that the Syrian hamster model was useful to detect differences in virulence of ancestral B.1 and the B.1.427 and B.1.429 epsilon VOCs by comparing body weight changes over 10 days. The timing and severity of lung pathology also distinguished the variants and correlated with the timing of virus dissemination into deep lung tissues based on ISH labelling. Further, the multi-virus in-vivo competition experiments provided insight into the relative fitness of the different SARS-CoV-2 variants and revealed that the anatomic site (lung vs URT) can affect the relative fitness of the variants. Clinical and epidemiologic studies are needed to confirm that the increased virulence and high relative fitness of the epsilon (427/429) variant in Syrian hamsters is mirrored in human infections.

## Materials and Methods

### Ethics statement

Approval of all animal experiments was obtained from the Institutional Animal Care and Use Committee of UC Davis, UC Berkeley, or the Rocky Mountain Laboratories. Experiments were performed following the guidelines and basic principles in the United States Public Health Service Policy on Humane Care and Use of Laboratory Animals and the Guide for the Care and Use of Laboratory Animals. Work with infectious SARS-CoV-2 strains under BSL3 conditions was approved by the Institutional Biosafety Committees (IBC). The male and female Syrian Hamsters used at UCD and UCB were 7-9 weeks old, and they were purchased from Charles River Inc. The male and female Syrian Hamsters used for the airborne transmission experiments at RML were 4-6 weeks old, and they were purchased from ENVIGO Inc. Inactivation and removal of samples from high containment was performed per IBC-approved standard operating procedures.

### Viruses and cells

SARS-CoV-2 variant strains ancestral B.1 (614G), B.1.427, B.1.429, B.1.1.7 were isolated from patient swabs by CDPH, Richmond CA, and P.2 and 351 were isolated from patient swabs at Stanford University, Stanford CA, USA. The “ancestral B.1 (614G)” SARS-CoV-2 variant had no defining mutations other than the B.1 (614G) mutation in spike. The P.1 variant was expanded by CDPH from a sample originally obtained from BEI Resources to produce the P1a stock. SARS-CoV-2 strain nCoV-WA1-2020 (lineage A) was provided by CDC, Atlanta, USA. Virus propagation was performed in Vero-86 cells in Dulbecco’s Modified Eagle Medium (DMEM) supplemented with 5% fetal bovine serum (FBS), 2 mM L-glutamine, 100 U/mL penicillin and 100 g/mL streptomycin. No contaminants were detected in any of the virus stocks except in the P.1 stock in which mycoplasma was detected. There is no indication that this contamination affected the experimental outcomes. All virus stocks except P1a were the second passage of the patient isolate on Vero cells, the original isolation being the first passage on Vero cells. The virus stocks were deep sequenced and the RNA sequences of all the virus stocks used were 100% identical to the corresponding variant sequences deposited in GenBank.

### Hamster inoculations

For experimental inoculations, 7-9 week old male Syrian hamsters (Charles River) were infected intranasally with a total dose of approximately 5000 PFU of SARS-CoV-2 suspended in 50μL sterile DMEM. For experiments in which mixtures of viruses were used, the dose of all variants was approximately equal (< 10-fold difference) and the total virus dose was approximately 5000 PFU. The airborne transmission experiments are described below.

### Infectious virus titer determination by TCID_50_ assay

Virus titer in lung was determined by a TCID_50_ assay in Vero E6 cells. Briefly, 10,000 cells per well were plated in 96-well plates and cultured for 24 hours at 37 °C/5% CO_2_. Lung tissue collected from hamsters was weighed and homogenized by bead beating. Homogenates were serial 10-fold diluted in Vero E6 growth medium and added to Vero E6 cells. Cells were observed for cytopathic effect for 5 days. TCID_50_ results were calculated using the Spearman and Kärber method (LOD 200 TCID_50_/mL) and normalized by lung tissue mass.

### qPCR for sub-genomic RNA quantitation

Quantitative real-time PCR assays were developed for detection of full-length genomic vRNA (gRNA), sub-genomic vRNA (sgRNA), and total vRNA. Upper respiratory tract washes and oral swabs were lysed in Trizol LS in BSL-3 and RNA was extracted from the aqueous phase in BSL-2. RNeasy mini kits (Qiagen) were used to purify the extracted RNA. Following DNAse treatment (ezDNAse; Invitrogen), complementary DNA was generated using superscript IV reverse transcriptase (ThermoFisher) in the presence of RNAseOUT (Invitrogen). A portion of this reaction was mixed with QuantiTect Probe PCR master mix and optimized concentrations of gene specific primers. All reactions were run on a Quantstudio 12K Flex real-time cycler (Applied Biosystems). sgRNA was quantified using primers sgLeadSARSCoV2_F (CGATCTCTTGTAGATCTGTTCTC) and wtN_R4 (GGTGAACCAAGACGCAGTAT), with probe wtN_P4 (/56-FAM/TAACCAGAA/ZEN/TGGAGAACGCAGTGGG/3IABkFQ/). Standard curves generated from PCR amplicons of the qPCR targets were used to establish line equations to determine RNA copies/mL of sample.

### Histopathology

At necropsy, lung was inflated with 10% buffered formalin (Thermo Fisher) and hamster tissues were fixed for 48 hours at room temperature in a 10-fold volume of 10% buffered formalin. Tissues were embedded in paraffin, thin-sectioned (4μm) and stained routinely with hematoxylin and eosin (H&E). H&E slides were scanned to 40x magnification by whole-slide image technique using an Aperio slide scanner with a magnification doubler and a resolution of 0.25 μm/pixel. Image files were uploaded on a Leica hosted web-based site and a board certified veterinary anatomic pathologist blindly evaluated sections for SARS-CoV-2 induced histologic lesions. For semi-quantitative assessment of lung inflammation, the pathologist estimated the area of inflamed tissue (visible to the naked eye at subgross magnification) as a percentage of the total surface area of the lung section. Each section of lung was further scored as described in Supplementary Table 1.

### In situ hybridization (ISH)

RNAscope^®^ ISH was performed according to the manufacturer’s protocol (Document Number 322452-USM and 322360-USM, ACD) with modifications. Briefly, we used RNAscope 2.5 HD Red Detection Kit (ACD) and RNAscope Probe - V-nCoV2019-S (cat# 848561, ACD) in the ISH assay. At each run of the ISH, RNAscope^®^ Negative Control Probe – DapB (cat# 310043) and tissues from a SARS-CoV-2 uninfected animal hybridized with the SARS-CoV-2 probes served as negative controls. RNAscope^®^ Probe - Mau-Ppib (cat# 890851, ACD) was used as a positive control for RNA quality and constancy of the ISH assay. Four-micron deparaffinized paraffin sections were pretreated with 1x Target Retrieval Buffer at 100^0^C for 15 minutes and RNAscope Protease Plus at 40^0^C for 30 minutes before hybridization at 40^0^C for 2 hours. A cascade of signal amplification was carried out after hybridization. The signal was detected using a Fast Red solution for 10 minutes at room temperature. Slides were counterstained with hematoxylin, dehydrated, cover-slipped, and visualized by using a bright field microscope.

### Quasispecies identification of low-frequency lineages of SARS-CoV-2 (QUILLS)

We designed a amplicon strategy named QUILLS (QUasispecies Identification of <u>Low-Frequency</u> Lineages of <u>SARS-CoV2) by which</u> all of the major circulating variants of concern (VOCs) and variants of interest (VOIs) could be identified on the basis of lineage-defining mutations in the SARS-CoV-2 spike and orf1b genes (Supplementary Figure 1). Five pairs of forward and reverse primers (Table 1, “QUILLS primer set”) were designed to span the nucleotide sites associated with the spike S13, W152, K417, L452, E484, N501, A570, D614, H665, P681, T1027, and V1176 and orf1b P976 and D1183 positions. Thus, these primers were designed to target mutations that would be able to discriminate between all VOCs and nearly all VOIs in a mixed population.

Extracted RNA was diluted 1:3, and 7 microliters of diluted material were used for first strand cDNA synthesis using ProtoScript^®^ II First Strand cDNA Synthesis Kit (New England Biolabs #E6560) with random hexamers and following manufacturer’s instructions. PCR amplification was performed using the QUILLS primer set and Q5^®^ High-Fidelity DNA Polymerase (NEB #M0491) as follows: 15.4 µl of water, 5 µl of Q5 reaction buffer, 0.5 µl of dNTPS (NEB N0447S), 0.25 µl of enzyme and 1.35 µl of primer for each pool (pool 1 and pool 2). Reactions were incubated at 98 °C for 30s, 35 cycles of 98 °C for 15s and 65 °C for 5m, 4 °C until use. Amplification product was bead washed with .8X CleanNGS Beads (CNGS-0500), DNA concentration was measured with qubit and all samples were normalized to 60 ng to avoid contamination. Libraries were prepared using NEBNext^®^ Ultra™ II DNA Library Prep Kit for Illumina^®^ (NEB #E7645L) following manufacturer’s instructions and NEB Next Multiplex Oligos for Illumina (96 Index Primers, NEB #E6609) for multiplexing. Final concentration was measured with qubit and final libraries were pooled for sequencing in a concentration between 20 and 40 ng. Final pool was diluted to 6.5 picomolar and spiked with 10% PhiX. MiSeq Reagent Kit v2 (300-cycles single-end) was used for sequencing on the Illumina MiSeq (Illumina MS-102-2002) following manufacturer’s specifications.

Raw FASTQ sequences were preprocessed using an in-house computational pipeline that is part of the SURPI software package [27,28]. The preprocessing step consisted of trimming low-quality and adapter sequences using cutadapt [29], retaining reads of trimmed length >75 bp, and then removing low-complexity sequences using the DUST algorithm in PrinSeq [30]. After filtering the preprocessed dataset for SARS-CoV-2-specific viral reads using the nucleotide BLAST algorithm with an e-value threshold cutoff of 10^-8^, lineage-specific mutation single nucleotide polymorphisms (SNPs) were identified and counted using a custom in-house computational script. The relative proportions of each SARS-CoV-2 variant in the mixture were estimated by manual analysis of the mutational frequencies at each of the lineage-defining SNPs.

### Airborne Transmission experiments

The airborne transmission experiments were performed at the Rocky Mountain Laboratories, NIAID, NIH as previously described [31]. Hamsters were co-housed (1:1) in specially designed cages with a perforated plastic divider dividing the living space in half, preventing direct contact between animals and movement of bedding material (alpha-dri bedding). Donor hamsters were infected intranasally with 10^4^ PFU of the different SARS-CoV-2 variants. Sentinel hamsters were placed on the other side of a divider 6-8 hours later. Sentinel hamsters received an oropharyngeal swab on 16 hours post exposure (PE), 24 hours PE, 2, 3, 5, 7, 10, and 14 days PE. Donors received an oropharyngeal swab 1 day post inoculation. Donors were euthanized at 7 or 8 days PI, sentinels were followed until 14 DPE. Experiments were performed with cages placed into a standard rodent cage rack, under normal airflow conditions. Sentinels were placed downstream of air flow.

Hamsters were weighed daily, and oropharyngeal (OP) swabs were taken daily until day 7 and then thrice a week. OP swabs were collected in 1 mL DMEM with 200 U/mL penicillin and 200 μg/mL streptomycin. Then, 140 μL was utilized for RNA extraction using the QIAamp Viral RNA Kit (Qiagen) using QIAcube HT automated system (Qiagen) according to the manufacturer’s instructions with an elution volume of 150 μL. Sub-genomic (sg) viral RNA and genomic (g) was detected by qRT-PCR [32,33]. Five μL RNA was tested with TaqManTM Fast Virus One-Step Master Mix (Applied Biosystems) using QuantStudio 6 Flex Real-Time PCR System (Applied Biosystems) according to instructions of the manufacturer. Ten-fold dilutions of SARS-CoV-2 standards with known copy numbers were used to construct a standard curve and calculate copy numbers/ml.

### Statistical analysis

Because the number of hamsters in each animal group was not uniform, mean values from each group were compared by fitting a mixed model, rather than by using a one-way ANOVA. A post-hoc multiple comparison test was used to compare the mean values of the B.1.427 or B.1.429 groups individually to the B.1 (614G) group. Graph Pad Prism 9.0 (San Diego, CA) installed on a MacBook Pro (Cupertino, CA) running Big Sur Version 11.5 was used for the analysis.

## Acknowledgements

We thank M. Ott, M. Busch, J. Derisi and D. Wadford and other members of the NorCal SARS-CoV-2 Assessment Group for helpful discussions and K. Keel for assistance with slide scanning. This work was supported by Intramural funding from the Center for Immunology and Infectious Diseases, UC Davis and NIH/R01-AI118590 to C.J. Miller; Fast Grants (part of Emergent Ventures at George Mason University) to S. Stanley; the Intramural Research Program of the National Institute of Allergy and Infectious Diseases, National Institutes of Health (1ZIAAI001179-01) to RML, a T32 AI007502-23 to A. Rustagi, NIH/R33-AI129455 and the US Centers for Disease Control and Prevention (contract 75D30121C10991) to C.Y. Chiu.

## Author Contributions

Conceptualization: CJM, CYC, SS, VM, CH, CAB

Data curation: CJM, CYC, TC, DF, NvD, LF, EB, MKM, AS-G, VS, AR, Z-MM

Formal analysis: CJM, CYC, TC, DF, NvD

Funding acquisition: CJM, CYC, SS, VM, CAB

Investigation: CJM, CYC, TC, DF, NvD, LF, EB, MKM, AS-G, VS, AR, CKY, JRP, Z-MM, MH, JS

Methodology: CJM, CYC, SS, VM

Project administration: CJM.

Supervision: CJM, CYC, SS, VM.

Writing – original draft: CJM

Writing – review & editing: CJM, CYC, SS, VM, CH, CAB, TC, DF, NvD, LF, EB, MKM, AS-G,VS, AR, CKY, JRP, Z-MM, MH, JS

## References

1. Otto SP, Day T, Arino J, Colijn C, Dushoff J, et al. (2021) The origins and potential future of SARS-CoV-2 variants of concern in the evolving COVID-19 pandemic. Curr Biol 31: R918–R929.

2. Gomez CE, Perdiguero B, Esteban M (2021) Emerging SARS-CoV-2 Variants and Impact in Global Vaccination Programs against SARS-CoV-2/COVID-19. Vaccines (Basel) 9.

3. Harvey WT, Carabelli AM, Jackson B, Gupta RK, Thomson EC, et al. (2021) SARS-CoV-2 variants, spike mutations and immune escape. Nat Rev Microbiol 19: 409–424.

4. Deng X, Garcia-Knight MA, Khalid MM, Servellita V, Wang C, et al. (2021) Transmission, infectivity, and neutralization of a spike L452R SARS-CoV-2 variant. Cell 184: 3426–3437 e3428.

5. Wu K, Werner AP, Koch M, Choi A, Narayanan E, et al. (2021) Serum Neutralizing Activity Elicited by mRNA-1273 Vaccine. N Engl J Med 384: 1468–1470.

6. McCallum M, Bassi J, De Marco A, Chen A, Walls AC, et al. (2021) SARS-CoV-2 immune evasion by the B.1.427/B.1.429 variant of concern. Science.

7. Boudewijns R, Thibaut HJ, Kaptein SJF, Li R, Vergote V, et al. (2020) STAT2 signaling restricts viral dissemination but drives severe pneumonia in SARS-CoV-2 infected hamsters. Nat Commun 11: 5838.

8. Munoz-Fontela C, Dowling WE, Funnell SGP, Gsell PS, Riveros-Balta AX, et al. (2020) Animal models for COVID-19. Nature 586: 509–515.

9. Chan JF, Zhang AJ, Yuan S, Poon VK, Chan CC, et al. (2020) Simulation of the Clinical and Pathological Manifestations of Coronavirus Disease 2019 (COVID-19) in a Golden Syrian Hamster Model: Implications for Disease Pathogenesis and Transmissibility. Clin Infect Dis 71: 2428–2446.

10. Imai M, Iwatsuki-Horimoto K, Hatta M, Loeber S, Halfmann PJ, et al. (2020) Syrian hamsters as a small animal model for SARS-CoV-2 infection and countermeasure development. Proc Natl Acad Sci U S A.

11. Zhou B, Thao TTN, Hoffmann D, Taddeo A, Ebert N, et al. (2021) SARS-CoV-2 spike D614G change enhances replication and transmission. Nature 592: 122–127.

12. Yinda CK, Port JR, Bushmaker T, Fischer RJ, Schulz JE, et al. (2021) Prior aerosol infection with lineage A SARS-CoV-2 variant protects hamsters from disease, but not reinfection with B.1.351 SARS-CoV-2 variant. Emerg Microbes Infect 10: 1284–1292.

13. Brocato RL, Principe LM, Kim RK, Zeng X, Williams JA, et al. (2020) Disruption of Adaptive Immunity Enhances Disease in SARS-CoV-2 Infected Syrian Hamsters. bioRxiv 20200619161612.

14. Carsana L, Sonzogni A, Nasr A, Rossi RS, Pellegrinelli A, et al. (2020) Pulmonary post-mortem findings in a series of COVID-19 cases from northern Italy: a two-centre descriptive study. Lancet Infect Dis 20: 1135–1140.

15. Mosleh W, Chen K, Pfau SE, Vashist A (2020) Endotheliitis and Endothelial Dysfunction in Patients with COVID-19: Its Role in Thrombosis and Adverse Outcomes. J Clin Med 9.

16. Ackermann M, Verleden SE, Kuehnel M, Haverich A, Welte T, et al. (2020) Pulmonary Vascular Endothelialitis, Thrombosis, and Angiogenesis in Covid-19. N Engl J Med 383: 120–128.

17. Calabrese F, Pezzuto F, Fortarezza F, Hofman P, Kern I, et al. (2020) Pulmonary pathology and COVID-19: lessons from autopsy. The experience of European Pulmonary Pathologists. Virchows Arch 477: 359–372.

18. Korber B, Fischer WM, Gnanakaran S, Yoon H, Theiler J, et al. (2020) Tracking Changes in SARS-CoV-2 Spike: Evidence that D614G Increases Infectivity of the COVID-19 Virus. Cell 182: 812–827 e819.

19. Faria NR, Mellan TA, Whittaker C, Claro IM, Candido DDS, et al. (2021) Genomics and epidemiology of the P.1 SARS-CoV-2 lineage in Manaus, Brazil. Science 372: 815–821.

20. Davies NG, Abbott S, Barnard RC, Jarvis CI, Kucharski AJ, et al. (2021) Estimated transmissibility and impact of SARS-CoV-2 lineage B.1.1.7 in England. Science 372.

21. Madhi SA, Baillie V, Cutland CL, Voysey M, Koen AL, et al. (2021) Efficacy of the ChAdOx1 nCoV-19 Covid-19 Vaccine against the B.1.351 Variant. N Engl J Med 384: 1885–1898.

22. Port JR, Kwe Yinda C, Avanzato VA, Schulz JE, Holbrook MG, et al. (2021) Increased aerosol transmission for B.1.1.7 (alpha variant) over lineage A variant of SARS-CoV-2. bioRxiv.

23. Teng S, Sobitan A, Rhoades R, Liu D, Tang Q (2021) Systemic effects of missense mutations on SARS-CoV-2 spike glycoprotein stability and receptor-binding affinity. Brief Bioinform 22: 1239–1253.

24. Chen J, Wang R, Wang M, Wei GW (2020) Mutations Strengthened SARS-CoV-2 Infectivity. J Mol Biol 432: 5212–5226.

25. Nunez IA, Lien CZ, Selvaraj P, Stauft CB, Liu S, et al. (2021) SARS-CoV-2 B.1.1.7 Infection of Syrian Hamster Does Not Cause More Severe Disease, and Naturally Acquired Immunity Confers Protection. mSphere: e0050721.

26. Abdelnabi R, Boudewijns R, Foo CS, Seldeslachts L, Sanchez-Felipe L, et al. (2021) Comparing infectivity and virulence of emerging SARS-CoV-2 variants in Syrian hamsters. EBioMedicine 68: 103403.

27. Naccache SN, Federman S, Veeraraghavan N, Zaharia M, Lee D, et al. (2014) A cloud-compatible bioinformatics pipeline for ultrarapid pathogen identification from next-generation sequencing of clinical samples. Genome Res 24: 1180–1192.

28. Miller S, Naccache SN, Samayoa E, Messacar K, Arevalo S, et al. (2019) Laboratory validation of a clinical metagenomic sequencing assay for pathogen detection in cerebrospinal fluid. Genome Res 29: 831–842.

29. Martin M (2011) Cutadapt removes adapter sequences from high-throughput sequencing reads. EMBnetjournal 17: 10–12.

30. Schmieder R, Edwards R (2011) Quality control and preprocessing of metagenomic datasets. Bioinformatics 27: 863–864.

31. Port JR, Yinda CK, Owusu IO, Holbrook M, Fischer R, et al. (2020) SARS-CoV-2 disease severity and transmission efficiency is increased for airborne but not fomite exposure in Syrian hamsters. bioRxiv.

32. Corman VM, Landt O, Kaiser M, Molenkamp R, Meijer A, et al. (2020) Detection of 2019 novel coronavirus (2019-nCoV) by real-time RT-PCR. Euro Surveill 25.

33. Wolfel R, Corman VM, Guggemos W, Seilmaier M, Zange S, et al. (2020) Virological assessment of hospitalized patients with COVID-2019. Nature 581: 465–469.

